# Valorization of mushroom by-products for sustainability: exploring antioxidant and prebiotic properties

**DOI:** 10.1101/2025.03.24.644873

**Authors:** Pablo Navarro-Simarro, Lourdes Gómez-Gómez, Ángela Rubio-Moraga, Elena Moreno-Gimenez, Alberto López-Jimenez, Alicia Prieto, Oussama Ahrazem

## Abstract

*Agaricus bisporus* (white button mushroom), *Pleurotus ostreatus* (oyster mushroom), and *Lentinula edodes* (shiitake mushroom) are the most cultivated mushrooms in the Spanish region of Castilla-La Mancha. The increasing production of these mushrooms has created opportunities for sustainable utilization of their by-products such as undersized mushrooms, stalks, and stems. This study evaluated the antioxidant, prebiotic, and antimicrobial properties of aqueous, ethanolic and alkaline extracts derived from these by-products. Aqueous extracts from oyster mushroom, rich in glucose-polysaccharides (59.4±0.4% mushroom dw), promoted the growth and lactic acid production of the probiotic bacteria *Lactocaseibacillus casei* and *Lactoplantibacillus plantarum*. Shiitake extracts demonstrated antimicrobial activity against *Pseudomonas aeruginosa*, *Escherichia coli* and *Salmonella enterica* (Minimal inhibitory concentrations: 15, 7.5 and 7.5 mg/mL, respectively), and stimulated the growth of lactic acid bacteria at low concentrations (1.875 mg/mL) but inhibiting them at higher concentrations. Extracts from white button mushrooms exhibited the strongest antioxidant activity, particularly ethanolic extracts rich in phenolic compounds (0.015 mg Gallic acid/mg extract). These results highlight the potential of extracts of mushroom by-products as organic sources of antioxidants, antimicrobials, and prebiotics, creating new avenues for food product development. Reusing these by-products could encourage sustainability and assist the mushroom sector in implementing zero-waste and circular economy methods.

## 1. Introduction

Given their nutritional significance and broad culinary applications, oyster mushrooms (*Pleurotus ostreatus*), button mushrooms (*Agaricus bisporus*) and shiitake mushrooms (*Lentinula edodes*) are among the most important crops grown worldwide [1]. However, this large-scale production generates significant by-products, such as stems and other mushroom wastes, which can account for up to half of the fruiting body mass. In Castilla-La Mancha, Spain, a well-known mushroom-producing region, these by-products, particulary those from *A. bisporus*, *P. ostreatus* stalks, and *L. edodes* stems, are mostly neglected. Despite their abundance and bioactive potential, the majority of these residues are consigned to low-value applications, such as animal feed, or are completely wasted, resulting in a missed opportunity for value generation [2,3].

Mushroom by-products have unexplored potential due to their rich composition, which includes polysaccharides, phenolic compounds, sterols, and various microelements [4–6]. Polysaccharides, such as β-glucans, are of particular interest due to their prebiotic characteristics. These compounds resists digestion and reach the gut microbiota, where they promote the growth of helpful bacteria such as *Lactocaseibacillus* spp. while suppressing the proliferation of pathogenic strains [7–11]. Furthermore, bioactive substances such as phenolic acids and peptides have strong antioxidant and antibacterial capabilities, which could provide health advantages and protect against oxidative stress and microbial infections [2,12–14].

Despite these promising attributes, mushroom by-products remain underexploited in both research and industry. Recent studies have begun exploring their incorporation into functional foods, leveraging their bioactive properties to enhance the nutritional and health-promoting profiles of various products. For instance, mushroom residues have been used to fortify meat and dairy products, enrich pasta, and serve as a natural flavor enhancer, replacing additives like monosodium glutamate [15–18]. However, the full potential of these residues, particularly their targeted application in prebiotic, antimicrobial, and antioxidant contexts, remains largely unexplored.

This study seeks to fill these gaps by focusing on the most troublesome by-products of *A. bisporus*, *P. ostreatus*, and *L. edodes* in Castilla-La Mancha. It assesses their prebiotic, antibacterial, and antioxidant activities using different extraction procedures, highlighting their potential for functional food creation. Unlike previous research, this study conducts a comprehensive assessment of region-specific by-products to investigate their bioactive potential with the dual goals of improving food systems and increasing sustainability. This work supports the circular economy’s precepts by converting low-value agricultural waste into high-value resources and demonstrating how bioactive extraction can solve environmental problems while also meeting consumer demand for functional foods. This dual approach highlights the potential for mushroom by-products to contribute meaningfully to sustainable food systems and human health, creating a model for zero-waste agricultural practices.

## 2. Materials and methods

### 2.1 Chemicals and reagents

Ethanol and sodium hydroxide were purchased from Honeywell. Gallic acid, triphenyl tetrazolium chloride, magnesium sulfate and manganese (II) sulfate were purchased from Sigma. Tween80, sodium acetate and yeast extract were purchased from Scharlab; sulfuric acid from Prolabo; Man Rogosa Sharpe broth (MRS) and Man Rogosa Sharpe Agar (MRSA) from Biokar; Folin and Ciocalteu’s phenol reagent and sodium carbonate from Panreac; 2,2-Diphenyl-1-Picrylhydrazyl from TCI; phenolic acid from Ambion and glucose, meat extract, European bacteriological agar and tryptone from Conda.

### 2.2 Mushroom by-products

The by-products (low-grade *A. bisporus*, *P. ostreatus* base, and *L. edodes* stem) were supplied by Setas Meli S.L., a company involved in the processing of edible mushrooms cultivated in Casasimarro (Albacete, Spain). These three species were selected due to their prominence in the Castilla-La Mancha region of Spain, where the disposal of these by-products poses a challenge for local businesses. The material was received within 12 h of harvesting and stored at-20°C, before being freeze-dried at-50°C for 48 h (LyoQuest-85 / 208 V 60 Hz, Teslar). Finally, the mushrooms were ground using a conventional mill, and the resulting powder was stored in hermetically sealed jars at room temperature. The three powders were designated as AB-Whole (*Agaricus Bisporus* – Whole Mushroom), PO-Stalk (*Pleurotus ostreatus* - Stalk), and LE-Stem (*Lentinula edodes* - Stem).

### 2.3 Extraction methods

The selection of extraction methods—aqueous, ethanolic, or alkaline—was chosen based on the chemical properties of the target bioactive compounds and their compatibility with the intended assays, ensuring optimal recovery and functionality. By aligning each method with the molecular characteristics of the bioactives and the specific requirements of the assays, this approach maximized the functional potential of mushroom by-products for applications in functional foods and nutraceuticals. These extraction protocols were designed with scalability in mind, ensuring their potential applicability in industrial settings. Aqueous, ethanolic, and alkaline extraction methods were selected for their simplicity, cost-effectiveness, and adaptability to larger-scale operations.

#### 2.3.1 Aqueous extracts

Aqueous extraction was employed to isolate hydrophilic compounds like polysaccharides, particularly β-glucans, which are critical for prebiotic activity [19]. For the preparation of aqueous extracts, 3 g of mushroom powder was added to 250 mL of distilled water and heated to 80°C with continuous magnetic stirring for 3 h [20]. Afterward, the mixture was then centrifuged at 8,000 rpm for 10 min to separate the insoluble fraction. The supernatant was then filtered using a 250 mL vacuum filter and stored under sterile conditions at 4°C, or frozen at-20°C for subsequent lyophilization. The aqueous extracts were labeled as AB-Whole-Aq, PO-Stalk-Aq, and LE-Stem-Aq.

#### 2.3.2 Ethanolic extracts

Ethanolic extraction, known for its efficiency in recovering phenolic compounds with strong antioxidant properties, was applied to assess radical scavenging and reducing capacity [21]. The ethanolic extract was prepared at a lower temperature than aqueous extracts to prevent the degradation of phenolic compounds and other antioxidants. Briefly, 3 g of mushroom powder was mixed with 100 mL of a 4:1 ethanol-water solution and incubated for 24 h at room temperature. Following incubation, the mixture was centrifuged, and the supernatant was concentrated using a rotary evaporator (Rotavapor R-210, Buchi, Flawil, Switzerland) at 40°C. After the ethanol had evaporated, the extract was lyophilized, and its yield was determined. The ethanolic extracts were designated as Ab-Whole-Et, PO-Stalk-Et, and LE-Stem-Et.

#### 2.3.2 Alkaline extraction

Alkaline extraction facilitated the release of complex and bound bioactives, such as polysaccharides (water-soluble and insoluble) and peptides, which are challenging to recover under neutral conditions. Briefly, 7 g of mushroom powder was mixed with 200 mL of 1 M NaOH and stirred magnetically for 3 h at room temperature. The mixture was then centrifuged at 10,000 rpm for 10 min to separate the supernatant. The pellet underwent a second extraction with another 200 mL of 1 M NaOH under the same conditions, followed by centrifugation. The supernatants from both extractions were combined, and absolute ethanol (v/v) was added to precipitate the polysaccharides. The resulting pellet was centrifuged and dialyzed against water using 10,000 Da cut-off membranes. Finally, the polysaccharide extract was lyophilized and stored for further use. The alkaline extracts were labeled as Ab-Whole-Alk, PO-Stalk-Alk, and LE-Stem-Alk.

### 2.4 Metabolite analysis

#### 2.4.1 HPLC analysis of the ethanol extracts

A Konik HPLC (Barcelona, Spain) equipped with a pump, autosampler, column oven, and an online photodiode array detector with a dynamic range of 200-700 nm was used to evaluate and quantify the extracts. For analytical purposes, Sugerlabor Inertsil ODS−2.5-µm C18 column (250×4.6 mm) was used. A procedure previously outlined for separation was followed [22]. The solvents used for sample elution were 90% acetonitrile with 0.05% TFA (solvent B) and 10% acetonitrile with 0.05% TFA (solvent A). The conditions were as follows: 0–5 min isocratic at 5% B, 5–40 min linear gradient from 5 to 25% B, 40–48 min linear gradient up to 100% B, 48–60 min isocratic at 100% of B and 60–72 min washing with 100% methanol at a flow rate of 0.2 mL/min. The system was then allowed to re-equilibrate for 10 min at initial conditions. The major compounds were identified using a homemade phenolic compound library and comparing the retention times and UV spectra of the peaks in the samples with those of authentic standards or data published in the literature [23].

#### 2.4.2 Total phenolic content

The total amount of phenols was expressed as gallic acid (GA) equivalents (mg GA/mg extract) as described by [24] with some modifications. Briefly, 5 µL of the sample was diluted in 495 µL of water and mixed with 250 µL of Folin and Ciocalteu’s phenol reagent. After 3 min, 1250 µL of Na_2_CO_3_ was added and the solution was allowed to stand for 30 min to obtain a bluish color, which was quantified at 760 nm using an iMark^TM^ Microplate Reader. A GA solution with concentrations ranging from 0.1 to 0.01 mg/mL was used as a standard.

#### 2.4.3 Total sugar analysis using phenol-sulfuric method

Total sugar content was calculated using phenol-sulfuric assay. For this purpose, 200 mL of phenol-chloroform (5:1) was added to 200 mL of the sample (1 mg/mL) and the reaction was incubated for 5 min at 70 °C. Next, 2 mL of H_2_SO_4_ was added to the solution, which was incubated for 10 min at room temperature, shaken, and incubated for another 30 min at room temperature. The absorbance of the solution was measured at 490 nm. Glucose (0.1-1 mg/mL) was used as a positive control. For the graphical comparison of the carbohydrate composition of the different extracts, the concentration of 1 mg/mL of glucose was used as 100% total sugar.

#### 2.4.4 Neutral sugar analysis by GC-MS

The analysis of neutral sugars was performed following the protocol described by [25]. The monosaccharide compositions of mushroom powder and polysaccharides were analyzed by GC-MS after acid hydrolysis (3 M trifluoroacetic acid, 1 h, 121 °C) and derivatization to trimethylsilyl oximes. Inositol was used as an internal standard.

### 2.5 Functional testing

#### 2.5.1 Bacterial strains

Bacterial strains were obtained from the Spanish Type Culture Collection (CECT). The probiotic bacteria *Lactocaseibacillus casei* CECT 475 and *Lactiplantibacillus plantarum* were selected as models for the study of the prebiotic potential of the extracts. These bacteria were chosen due to their well-documented roles as probiotics in both industrial and health contexts. These strains are widely used in functional food formulations and supplements for their ability to enhance gut health, boost the immune system, and inhibit pathogenic bacteria through competitive exclusion and lactic acid production. Their relevance extends to diverse applications, including fermented dairy products, plant-based beverages, and gut microbiome-targeted therapies, underscoring their industrial and therapeutic importance [26]. In addition, pathogenic strains of *Salmonella enterica*, *Escherichia coli* and *Pseudomonas aeruginosa* were used for antimicrobial assays, as these bacteria are commonly associated with food spoilage [27]. All bacteria were grown at 37°C for 24 h in optimal media (MRSA for probiotic strains and Lysogeny Broth agar (LBA) for pathogenic strains). All strains were stored at 4°C and subcultured bimonthly to ensure viability.

To standardize the bacterial concentrations before the assays, the bacteria were grown in liquid medium at a concentration of 10^8^ CFU/mL. For this analysis, their OD was measured at 600 nm using an iMark^TM^ Microplate Reader [28].

#### 2.5.2 2,2-Diphenyl-1-Picrylhydrazyl (DPPH) radical scavenging activity

The antioxidant potential of the extracts was elucidated by the DPPH method using a solution of the radical at 200 µM and serial dilutions of the extracts to be screened (2 - 0.063 mg/mL). Briefly, 150 µL of the DPPH solution was mixed with 50 µL of the diluted extract, and the absorbance was measured at 517 nm immediately (A0) and after 30 min (A1). The percentage of free radicals inhibition was measured using the following formula: % = (A0-A1/A0)x100 [29]. Finally, distilled water was used as negative control and ascorbic acid 0,01 M as positive control for this assay.

#### 2.5.3 Microdilution method for antimicrobial analysis

Overnight cultures of bacteria were grown and normalized to a concentration of 10^8^ CFU/mL. Following the protocol of [30], serial dilutions of the extracts (15 mg/mL - 1.88 mg/mL) were performed in a volume of 100 µL per well. In addition to the dilutions, 100 µL of bacteria were inoculated to obtain a final volume of 200 µL. The 96-well plates were incubated with lid for 24 h at 37°C and the turbidity of the wells was measured at 600 nm. To confirm the minimum inhibitory concentration (MIC), 10 µL of a 0.5% solution of triphenyl tetrazolium chloride was added.

#### 2.5.4 Prebiotic potential analysis

##### 2.5.4.1 Formulation of broth for probiotic bacteria

The evaluation of prebiotic potential used *L. casei* and *L. plantarum* as model probiotic bacteria. Low-glucose MRS and minimal medium media were formulated based on commercial MRS (Table 1). Acetic acid was added to obtain a pH of 6, and the media were filtered and stored in sterile conditions until use.

**Table 1.** Formulation of commercial MRS, low glucose MRS and minimal medium for *L. casei* and *L. plantarum*.

| Reagents | Commercial MRS<br>(g/L) | Low glucose MRS<br>(g/L) | Minimal medium<br>(g/L) |
| --- | --- | --- | --- |
| Glucose | 20 | 2 | - |
| Peptone | 20 | 1 | - |
| Yeast extract | 5 | 0.40 | - |
| Dipotassium phosphate | 2 | 1 | 1 |
| Sodium acetate | 5 | 0.75 | 0.75 |
| Magnesium sulfate | 0.2 | 0.1 | 0.1 |
| Manganese sulfate | 0.05 | 0.025 | 0.025 |
| Tween80 | 1.08 | 0.5 (mL) | 0.5 (mL) |

##### 2.5.4.2 High-volume fermentation

To test the prebiotic potential of the aqueous extracts, 3 g of mushroom powder were extracted with 250 mL of minimal medium for 3 h at 80°C. The extracts were filtered and stored until use. The fermentation assays were performed in 100 mL Erlenmeyer flasks containing 60 mL of the previous extract and inoculated with 1 mL of the bacterial culture at a concentration of 10^8^ CFU/mL. The flasks were sealed to minimize oxygen exposure and were incubated at 37°C for 24 h with shaking. Subsequently, samples were collected to measure bacterial growth at 600 nm and lactic acid concentrations using the protocol of [31]. As controls, MRS and inulin (2 g/L) were used.

##### 2.5.4.3 Growth evaluation by microdilution method

For the 96-well plate assays, the previously described microdilution method was followed and bacterial growth was recorded at 24 h. The assays were performed using minimal medium and low glucose medium supplemented with different concentrations of each aqueous extract (0.234 - 15 mg/mL). For comparison, bacterial growth was evaluated in two control conditions: minimal or low glucose medium with increasing concentrations of inulin (0.234–15 mg/mL), and the other using low glucose MRS medium alone as a positive control.

### 2.6 Statistical analysis

Experiments were performed in triplicate and are presented as: mean ± standard deviation. The GraphPad Prism 8.0.2 statistical package was used for data analysis using one-way ANOVA tests followed by Tukey’s and Dunnett’s post hoc tests to compare means. Differences were considered significant when *p* < 0.05.

## 3. Results

### 3.1 Dehydration methods

Significant differences (*p* < 0.05) were observed between the water content of the three mushroom by-products, evaluated by freeze-drying and hot air drying. Among them, Ab-Whole possessed the highest amount of water, followed by PO-Stalk and LE-Stem. There were no significant differences between the values determined by the two treatments for any of the by-products tested (Figure 1A). Despite this, the samples treated with hot air drying showed a strong browning, which indicates degradation of their bioactive compounds. Therefore, to obtain a powder as close as possible to the fresh by-product, the lyophilized products were selected as the starting point for the preparation of the extracts.

**Figure 1.**
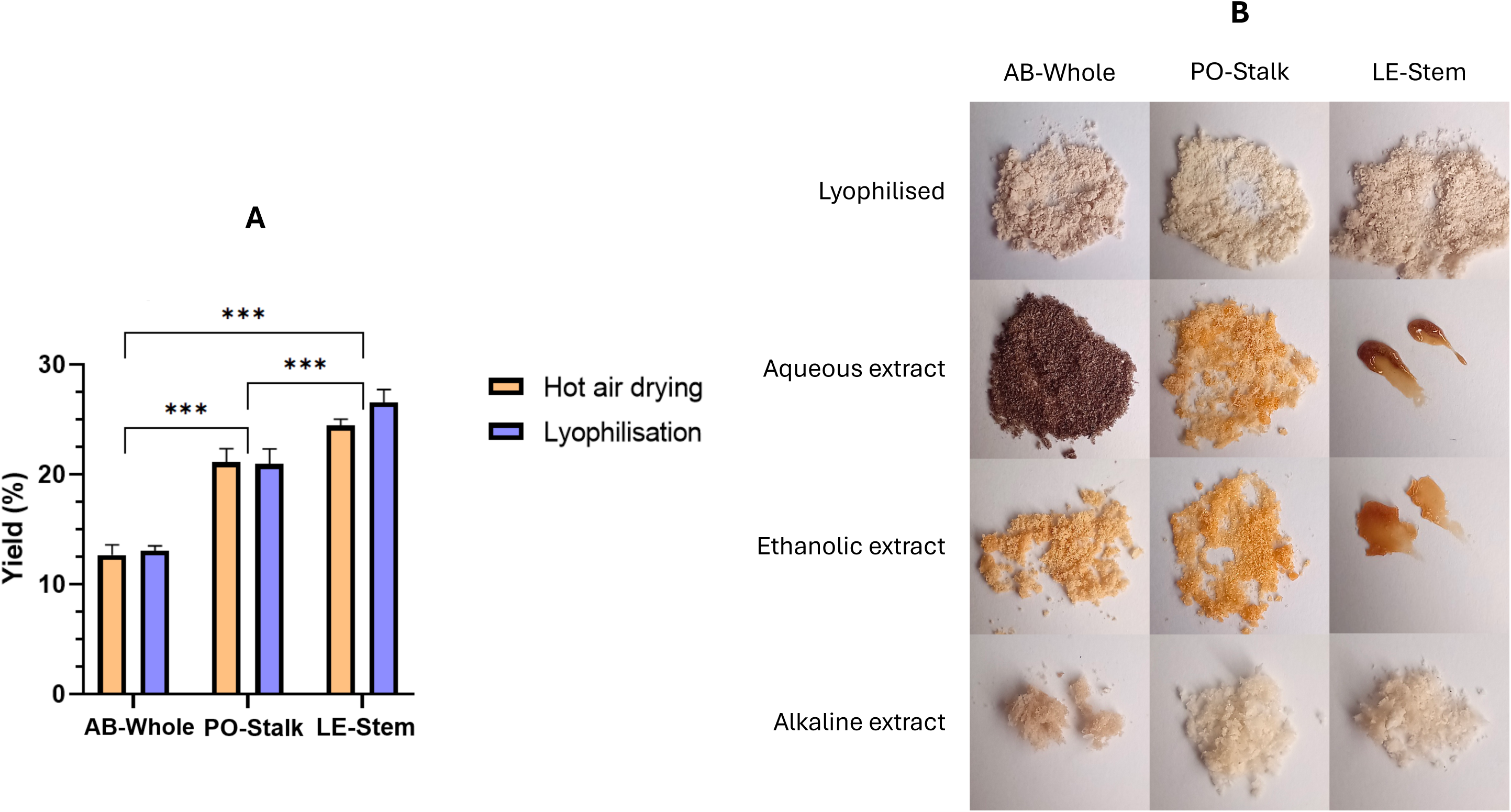
Comparison of the different dehydration and extraction methods. (A) Yields (%) obtained for each of the three mushroom by-products when dehydrated using either the hot air drying or lyophilisation method. Error bars represent the average values ± SD (n = 3). Asterisks denote significant differences between yields of each by-product (**** p < 0.0001); (B) Visual appearance of the different mushroom powders and extracts.

### 3.2 Extraction methods

The fungal by-products exhibited differences with respect to the extraction methods. For the aqueous extracts, the highest yield was obtained using Ab-Whole-Aq (56.3±0.8%), followed by PO-Stalk-Aq and LE-Stem-Aq (35.0±1.0% - 34.6±0.5%). The most differential characteristic of Ab-Whole-Aq was its dark color compared with the other extracts (Figure 1B), which could be due to the action of polyphenol oxidases, tyrosinases and laccases which are released when fungal membranes are disrupted [32]. This darkening was not appreciable in Ab-Whole-Et (25.1±0.6%) or in the remaining ethanolic extracts (PO-Stalk-Et: 19.0±0.5%; LE-Stem-Et: 12.1±0.3%). On the other hand, the alkaline extracts were characterized by the insolubility of most of the extracted compounds in water, since this method recovers proteins and both soluble and insoluble polysaccharides [33]. The extraction yields were 17.9±0.3%, 57.3±0.9% and 47.3±1.0% for Ab-Whole-Alk, PO-Stalk-Alk and LE-Stem-Alk, respectively.

### 3.3 Major bioactive compounds in the ethanolic extracts

The main compounds in the ethanolic extracts were analyzed by HPLC-DAD. In Ab-Whole-Et, gallic acid and caffeic acid were detected, in addition to other phenolic compounds that could not be identified due to their low concentrations (Figure 2C). Both gallic acid and caffeic acid are known for their antioxidant potential [34], which has been corroborated by DPPH assay. Regarding PO-Stalk-Et, its phenolic concentration was low and no compounds could be identified by HPLC-DAD. Finally, in the LE-Stem-Et analysis, a peak corresponding to eritadenine was clearly detected. This secondary metabolite present in *L. edodes* has been studied for its anxiety regulation effects [35].

**Figure 2.**
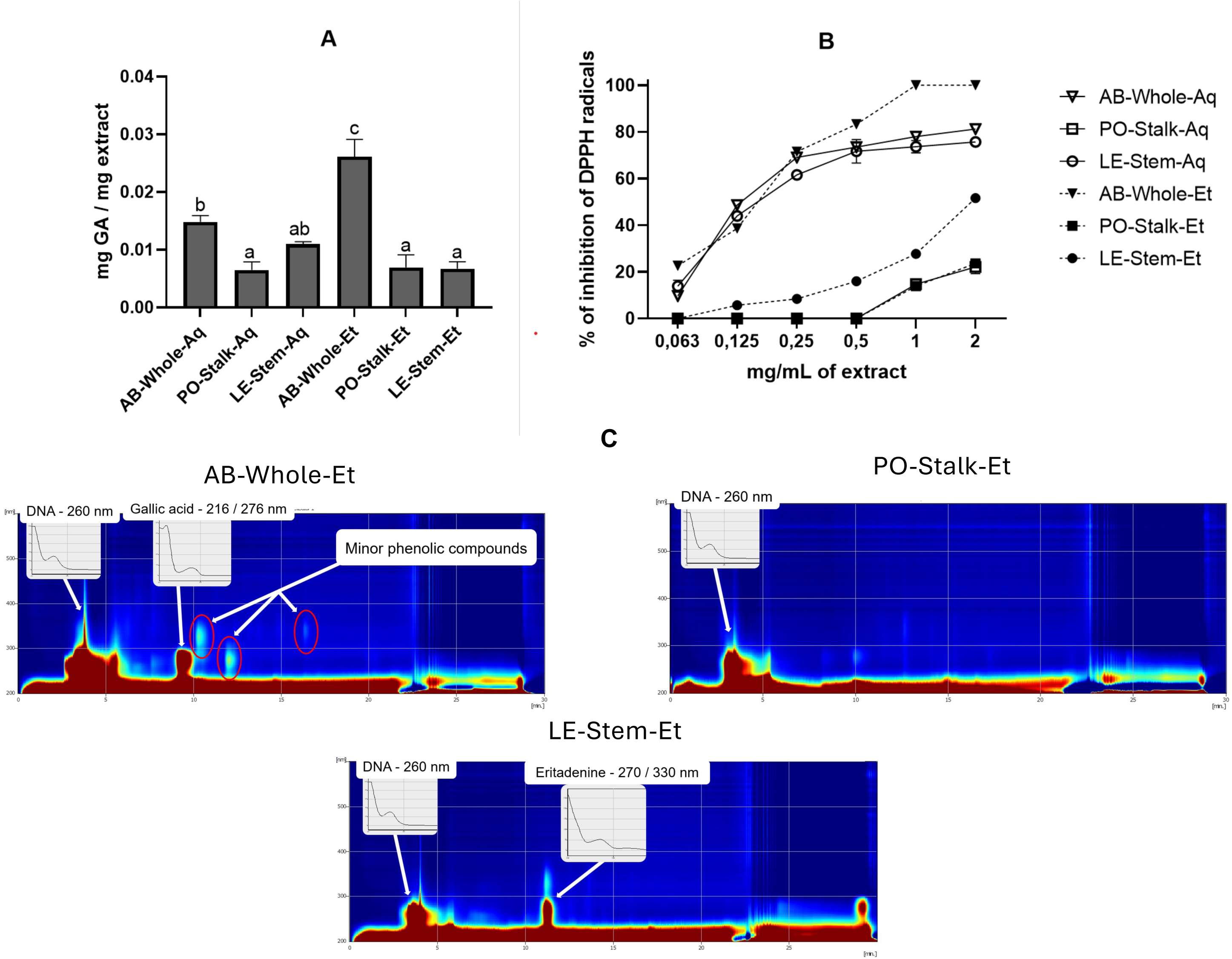
Antioxidant potential of mushroom extracts. (A) Results of the Folin-Ciocalteu assay. The graph represents the phenolic compound content of the extracts expressed as mg of gallic acid per mg of extract. Statistical analysis was performed using Tukey’s test, with letters (a, b, c) indicating significant differences between the samples; (B) The graph shows the results of the DPPH assay, where the antioxidant potential of the extracts is compared. The percentage of radical inhibition against increasing concentrations of the extract is evaluated; (C) Spectra show retention time (min) versus absorbance (nm) for Ab-Whole-Et, PO-Stalk-Et and LE-Stem-Et extract and the identification of major metabolites.

### 3.4 Phenolic content and antioxidant activity of the aqueous and ethanolic extracts

The phenolic compounds present in the fungal extracts specifically reacted with the Folin-Ciocalteu reagent to form a blue complex that is quantifiable in the visible spectrum. Significant differences were observed in the total phenolic content among the different by-products, as well as between the aqueous and ethanolic extracts of Ab-Whole and LE-Stem (Figure 2A). The alkaline extracts were omitted from Figure 2A due to their low concentration of phenolic compounds. The whole by-products and their alkaline extracts were omitted from this assay, because of the large number of insoluble compounds that interfered with the quantification. It should be noted that the ethanolic and aqueous extracts recovered from Ab-Whole, the by-product that contains the whole *A. bisporus* mushroom, presented the highest concentrations of antioxidant compounds. This was in agreement with previous studies that have provided evidence supporting the notion that mushroom caps serve as reservoirs for bioactive compounds, whereas the stems play a role in transporting these compounds throughout the organism [36].

These results were reinforced by the DPPH assay, in which the antioxidant potential of these extracts was compared (Figure 1B). Due to the lack of linearity of this assay at very high or low concentrations, the percentage inhibition was expressed as greater than 90% or less than 5%. The Ab-Whole-Et extract displayed a very strong antioxidant capacity (>90%) at concentrations ≤ 1 mg/mL, and maintained values of over 70% at concentrations up to 0.25 mg/mL. The antioxidant potential was immediately followed by the Ab-Whole-Aq extract and, in fact, the inhibition values started to equal those observed for Ab-Whole-Et at concentrations of approximately 0.25 mg/mL. LE-Stem-Aq also showed strong antioxidant activity (over 70%) at the highest concentrations tested, while the capacity of the LE-Stem-Et extract was much lower and both PO-Stalk extracts exhibited poor antioxidant properties at any concentration evaluated. These findings revealed a clear correlation between antioxidant potential and total phenol content determined by the Folin-Ciocalteu assay.

### 3.5 Antimicrobial effect

Among all the extracts tested, only LE-Stem-Aq and Ab-Whole-Et had an effect against the pathogenic bacteria tested at concentrations equal to or less than 15 mg/mL of extract. Between them, the shiitake extract had the lowest MIC, being 15 mg/mL for *P. aeruginosa* and *E. coli* and 7.5 mg/mL for *S. enterica* (Figure 4). It has been described that numerous extracts of *L. edodes* have antimicrobial activity against periodontal pathogens [37] as well as the effect of aqueous extracts of some of its varieties that possess an effect against antibiotic-resistant bacteria such as methicillin-resistant *S. aureus* [38].

### 3.6 Total sugar analysis

The phenol-sulfuric acid method is a traditional method for quantifying total sugars contents in biological samples. A key advantage of this method is its immediacy. However, it is important to indicate that certain non-glycosidic molecules can react with acids to produce colored compounds [39]. In addition, the calibration curve was generated using glucose, but the color intensity can vary for other monosaccharides, and the presence of natural phosphorylated polysaccharides in fungi could also be a source of variability in this assay [39]. Thus, the values estimated in this assay, as represented in Figure 3, could be under-or overestimated. In spite of this, the results are useful for comparing the total amount of carbohydrates in the different by-products and their extracts. Among them, it was observed that PO-Stalk and its alkaline, ethanolic and aqueous extracts had the highest carbohydrate content compared to analogous materials from LE-Stem and Ab-Whole. The presence of carbohydrates was notably lower in the aqueous and ethanolic extracts, except for PO-Stalk-Et, compared to the alkaline extracts, which present a large part of the soluble and insoluble polysaccharide fraction of the fungus.

**Figure 3.**
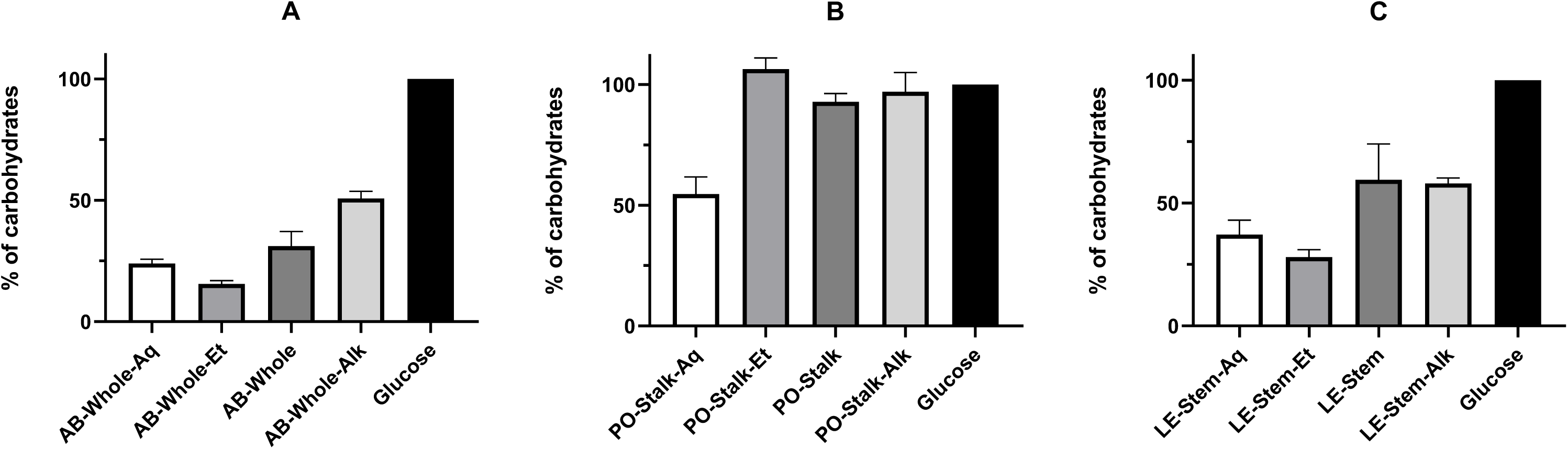
Sugar content of the different extracts. Percentage of sugars of AB-Whole extracts (A), PO-Stalk extracts (B) and LE-Stem extracts (C), relative to the control in the phenol-sulfuric assay. Error bars represent the average values ± SD (n = 3). Asterisks denote significant differences between yields obtained for each by-product (**** p < 0.0001).

**Figure 4.**
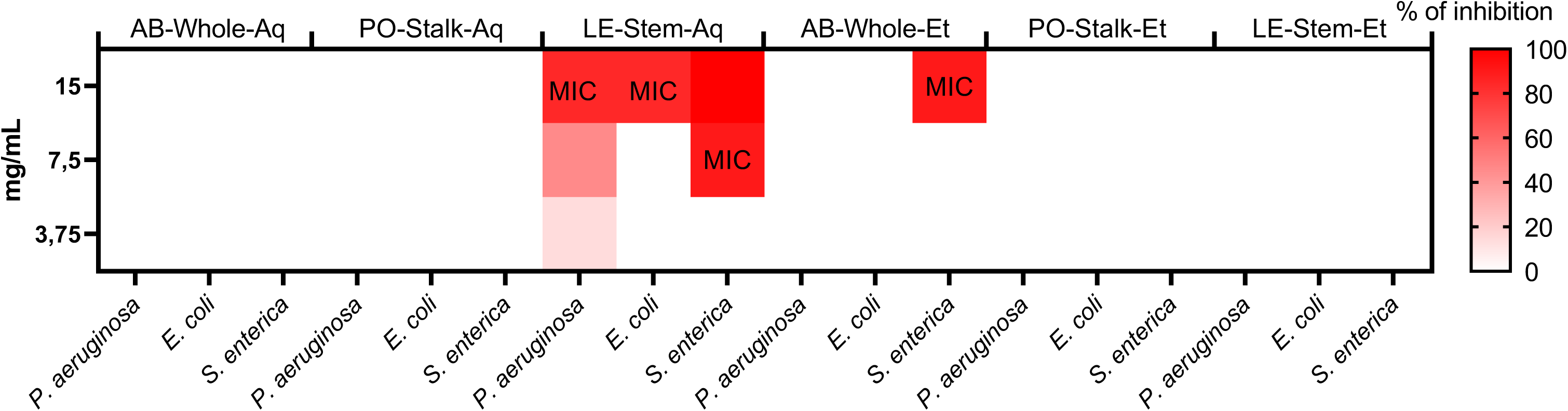
Antimicrobial effects of mushroom extracts. Percentage of growth inhibition of *P. aeruginosa*, *E. coli* and *S. enterica* against different fungal extracts. The minimum inhibitory concentration (MIC) of each extract is indicated within a range of 15 to 3.75 mg/ml.

GC-MS was used to identify the monosaccharides present in the by-products and their extracts to obtain a more accurate concentration of total carbohydrates. Glucose was identified as the main component in all samples, but quantitative differences were observed, since PO-Stalk and LE-Stem contained much more sugar than Ab-Whole (Table 2). Alkaline extraction is often preferred for its ability to break down cell walls, thus allowing for the recovery of complex polysaccharides like chitin and glucans, which may be otherwise inaccessible through aqueous or ethanolic extractions. Taking into account the percentage of glucosamine, it is also possible to estimate the presence of chitin in all samples, which is a highly valuable polysaccharide since its deacetylation gives rise to chitosan, which has in turn multiple applications [29,40]. Among the aqueous and ethanolic extracts, those derived from Ab-Whole had the lowest proportion of monosaccharides. In contrast, extracts formulated from mushroom stems, LE-Stem and PO-Stalk, exhibited a higher percentage of sugars. The PO-Stalk-Aq extract, containing 31.4 ± 0.5% monosaccharides by dry weight, far exceeds the sugar content of Ab-Whole-Aq and Ab-Whole-Et extracts. This significant enrichment suggests that PO-Stalk-Aq could serve as a more effective prebiotic substrate due to its ability to provide a more diverse and abundant source of carbohydrates for bacterial fermentation.

**Table 2.** Monosaccharides identified by GC-MS of alkaline extracts and mushroom powder by GC-MS.

| Sample | Monosaccharide (% of dry weight) |  |  |  | Other compounds |
| --- | --- | --- | --- | --- | --- |
|  | Glucose | Galactose | Mannose | Glucosamine |  |
| Ab-Whole | 9.2±0.2 | 0.8±0.1 | 0.6±0.1 | 1.6±0.1 | - |
| Ab-Whole-Aq | 2.1±0.1 | - | 1.4±0.2 | - | Mannitol |
| Ab-Whole-Et | 2.1±0.1 | - | - | - | Mannitol |
| Ab-Whole-Alk | 39.7±0.4 | 1.7±0.2 | 0.5±0.1 | - | - |
| PO-Stalk | 59.4±0.4 | 0.7±0.1 | 1.6±0.1 | 0.8±0.1 | - |
| PO-Stalk-Aq | 31.4±0.5 | 3.1±0.1 | 2.1±0.2 | - | Mannitol |
| PO-Stalk-Et | 35.6±0.2 | - | - | - | Mannitol |
| PO-Stalk-Alk | 68.9±0.4 | - | 1.0±0.1 | 0.8±0.1 | - |
| LE-Stem | 45.4±0.3 | 0.8±0.1 | 2.6±0.1 | 0.8±0.1 | Xylitol, mannitol, and glucitol |
| LE-Stem-Aq | 10.7±0.2 | 2.7±0.2 | 1.6±0.2 | - | Mannitol |
| LE-Stem-Et | 8.6±0.1 | - | - | - | Mannitol |
| LE-Stem-Alk | 75.5±0.6 | 0.7±0.1 | 1.8±0.1 | 1.0±0.1 | - |

### 3.6 Prebiotic potential

*Lactobacillus* and its associated genera comprise bacteria with major importance in the balance of the human microbiome [41,42]. In various industries, some strains are used in the fermentation of certain foods such as yogurts and meat products [43], or to produce lactic acid [44]. The aim of this experiment was to compare the viability, growth, and metabolic activity of *L. plantarum* and *L. casei* in minimal medium supplemented with aqueous extracts of *A. bisporus*, *P. ostreatus* and *L. edodes* or with inulin, a well-known prebiotic polysaccharide, and in MRS medium. Numerous studies have analysed the effect of supplementing MRS with fungal extracts [45], although few have investigated the supplementation of a minimal medium [46]. The choice of aqueous extracts to evaluate their prebiotic effects is based on three factors: the safety of the solvent for use in food, its low cost and the evidence of the presence of prebiotic β-glucans by this extraction method [45].

Data obtained from microplate assays in minimal medium supplemented with the three extracts indicated that Ab-Whole-Aq and PO-Stalk-Aq were able to maintain bacterial viability and even promoted the growth of both strains without the need for additional nutrients (Figures 5A and 5B). In contrast, the use of LE-Stem-Aq slow down or inhibited bacterial growth at any dose. This was evident from the results obtained in the cultures additionally supplemented with glucose (Figures 5C and 5D), where the growth of both microorganisms was slightly inhibited by high concentrations of the LE-Stem-Aq extract. In contrast, at medium concentrations (1.875 mg/mL), the bacteria even increased their growth. These differences could be due to the presence of other macro-and microelements in Ab-Whole-Aq and PO-Stalk-Aq that are necessary for the correct development of the microorganisms in addition to sugars, or to the presence of some growth inhibitors in LE-Stem-Aq. Additionally, both bacteria exhibited reduced or no growth when supplemented with the inulin extract, possibly due to the absence of a nitrogen source.

**Figure 5.**
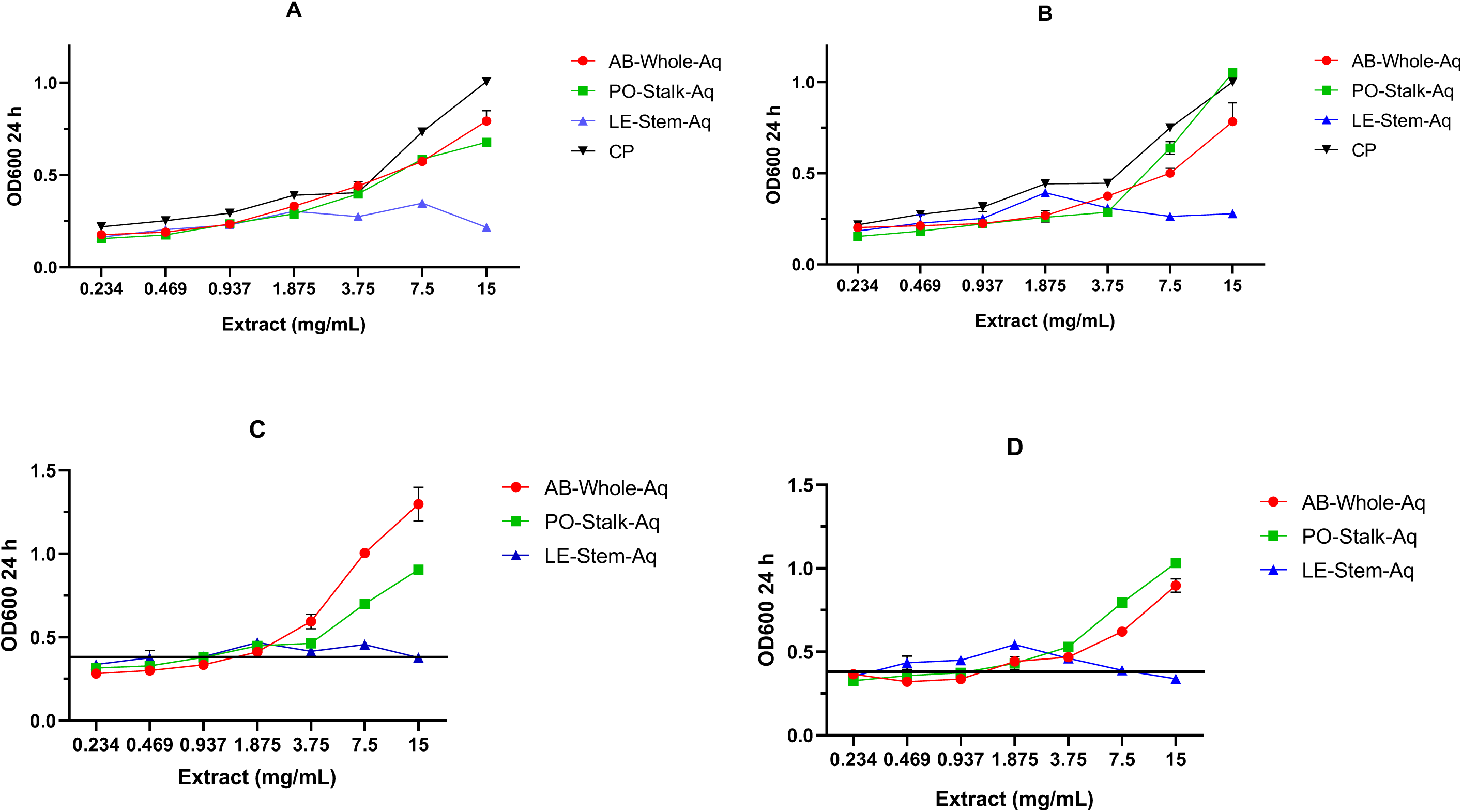
**Prebiotic effects of the extracts on *L. casei* and *L. plantarum* model bacteria by microdilution method**. (A) *L. casei* without glucose, microdilution method; (B) *L. plantarum* without glucose, microdilution method; (C) *L. casei* + glucose, microdilution method; (D) *L. plantarum* + glucose, microdilution method; The black line marked in Figures 4C and 4D refers to the mean bacterial growth of the culture supplemented only with MRS medium.

The inhibitory effects observed with higher concentrations of LE-Stem-Aq on the growth of both *L. plantarum* and *L. casei* could be attributed to several factors. First, the higher concentrations of LE-Stem-Aq may contain growth inhibitors or compounds that are not conducive to bacterial growth. These compounds could potentially interfere with the bacteria’s metabolism or cell membrane integrity, leading to reduced viability or inhibited growth. Additionally, the presence of some bioactive compounds, such as phenolic acids or secondary metabolites, at higher concentrations may exhibit antimicrobial properties that limit the growth of both the probiotic strains. It is also possible that, at higher concentrations, the osmotic stress caused by increased solute concentrations in the medium could impact bacterial growth. In contrast, at lower concentrations, LE-Stem-Aq may provide a more balanced environment, allowing the bacteria to grow and thrive by supplying essential nutrients while avoiding the inhibitory effects seen at higher concentrations. This suggests a concentration-dependent effect of LE-Stem-Aq, where a moderate concentration can support the metabolic activity and growth of the probiotics, while higher concentrations may have a negative impact.

In the fermentation assays performed with 60 mL of medium, the results varied with respect to those obtained in the microdilution trials due to the different yields of each extraction. Under these conditions, bacterial growth was significantly higher (**** p < 0.0001) using minimal medium with PO-Stalk-Aq for both strains (Figure 6A). Oyster mushroom extract remained above the control, and Ab-Whole-Aq exhibited a similar growth stimulation to control in *L. casei* culture. As observed in the microdilution assays, there was no bacterial growth in either the LE-Stem-Aq-supplemented or inulin-supplemented medium. Consistent with these results, a much higher lactic acid concentration was detected when bacteria were supplemented with PO-Stalk-Aq (Figure 6B). Since the amount of lactic acid produced directly reflects the metabolic activity of the bacteria, this finding points out that PO-Stalk-Aq is an excellent substrate for the growth and activity of both bacteria.

**Figure 6.**
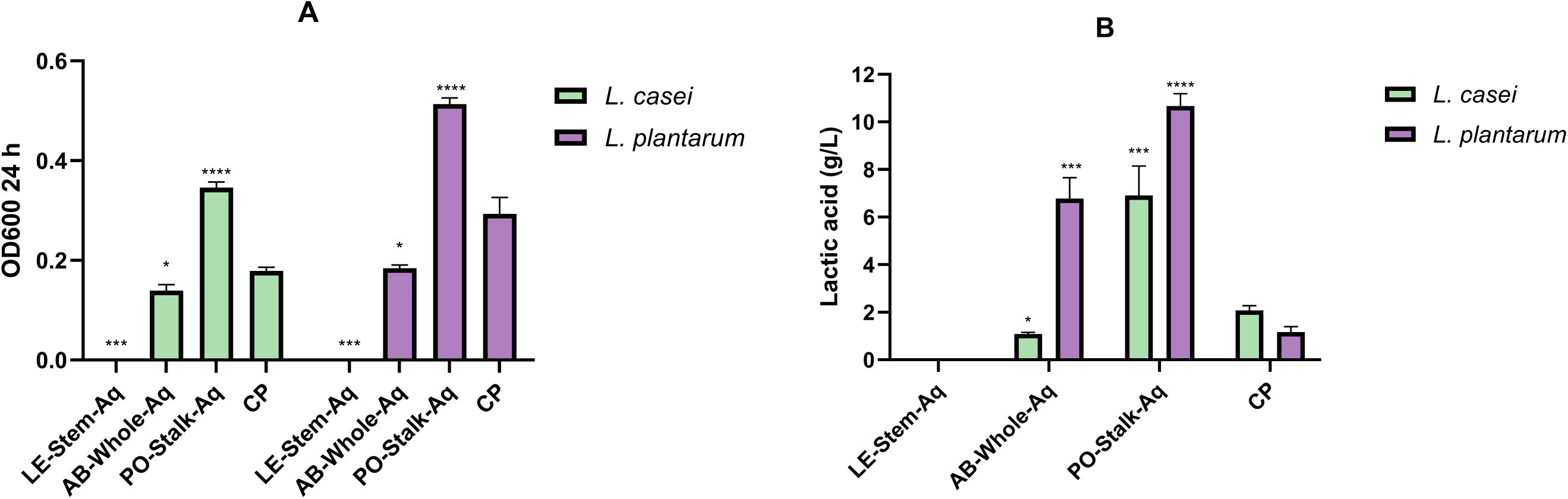
Prebiotic effects of fungal extracts on probiotic bacteria in large-volume fermentations. (A) Bacterial growth (OD600 24 h) for each strain grown for 24h in 60 mL of minimal medium; (B) Lactic acid concentration (g/L) of bacterial cultures grown for 24h in 60 mL of minimal medium. In all figures, positive control (CP) corresponds to the medium supplemented with glucose.

## 4. Discussion

Mushroom farming yields large amounts of by-products, such as spent mushroom substrate, stalks and non-commercial mushrooms. These by-products, which were once regarded as waste, are now receiving a lot of interest and attention in the scientific and industrial communities because of their potential as a source of bioactive compounds for the pharmaceutical and food industries [3].

Several studies have demonstrated the antimicrobial properties of mushroom extracts against a variety of microorganisms, including fungi, yeasts, and bacteria [47,48]. Bioactive substances such as polysaccharides, peptides, and phenolic compounds are thought to be responsible for the antimicrobial potential of mushroom extracts [37]. For *L. edodes* and *A. bisporus*, the antimicrobial potential of ethanolic extracts from the “cap” has been demonstrated, with MICs ranging from 5 to 10 mg/mL against gram-positive and gram-negative pathogenic bacteria [12]. Phenolic compounds and other secondary metabolites would act by affecting the permeability of the bacterial membrane, a mechanism similar to that of phenolic extracts derived from fruit by-products [28]. This study further shows that shiitake stems extracts also have antimicrobial activity, although with a higher MIC range.

The *in vitro* results of the prebiotic effect of PO-Stalk-Aq are in agreement with studies [49] that reported an increase in the growth or metabolic activity of *Lactobacillus* strains when supplementing MRS medium with the extract of this fungus. In our comparison of the prebiotic effects of fungal extracts with and without exogenous glucose, both Ab-Whole-Aq and PO-Stalk-Aq supported bacterial survival and growth in minimal media. However, at higher broth volumes, Ab-Whole-Aq’s prebiotic effect diminished considerably. In contrast, the sugars detected by GC-MS in PO-Stalk-Aq suggest that its higher carbohydrate content makes it an optimal substrate for these bacteria. This prebiotic effect has been demonstrated in aqueous extracts from *Pleurotus*, which have a total polysaccharide content of 50-70%, of which 30-50% are β-glucans [11]. These polysaccharides are composed of a chain of glucoses (1→3) and branches (1→6), which in GC-MS is seen as 59.6% glucoses in PO-Stalk (free and part of the polysaccharide structure), which resembles the proportion of other extracts made with the whole fungus.

The prebiotic results for the shiitake extracts are slightly different from those of previous studies. Although the antimicrobial effects of some of these compounds have already been demonstrated [37,50], their prebiotic effects were not significant under the conditions assayed [45]. This is because, our study distinguishes itself from many studies that focus solely on specific polysaccharide fractions by evaluating the total extract, thereby capturing the synergistic effects of the entire polysaccharide fraction. Despite this, as seen in prebiotic assays using microdilutions, LE-Stem-Aq slightly promoted the growth of both *Lactobacillus* strains at a concentration of 1.875 mg/mL, whereas higher concentrations resulted in some inhibition which leading to an OD_600_ matching the control in glucose-supplemented assays. In the food field, this behavior is not undesirable behavior, since a high concentration of LE-Stem-Aq as an antimicrobial and reinforces the umami taste of the product, as reported in fermented meat products, such as sausages [51]. Notably, antimicrobial assays on gram-negative pathogenic bacteria indicate that shiitake extracts may achieve complete inhibition of these pathogens, while allowing probiotic bacteria to retain their activity. This specific antibacterial activity highlights shiitake extracts’ potential as powerful prebiotic candidates [52]. Finding a suitable range of applications for this by-product is a new challenge, although many studies have already used it for food formulation [53].

Regarding antioxidant potential, *A. bisporus* showed a higher capacity with both Ab-Whole-Aq and Ab-Whole-Et, with Ab-Whole-Et being the most potent extract. In the HPLC analysis, a high amount of gallic acid was also observed in Ab-Whole-Et, which is in agreement with other studies attributing the antioxidant effect of ethanolic mushroom extracts to phenols [54,55]. Regarding its use in the food industry, Ab-Whole-Aq would have a more limited market because it produces a dark brown color in food due to its susceptibility to enzymatic browning [32]. The remaining extracts, except LE-Stem-Aq, showed much lower antioxidant capacity and total phenolic content. This was because the extracts of *A. bisporus* were obtained from the fruiting body of the mushroom, whereas the extracts of *P. ostreatus* and *L. edodes* were obtained from the stems. In mushrooms, stems function as transporters of biomolecules, while the caps act as reservoirs of these compounds [36].

The increasing global demand for edible mushrooms is expected to generate large quantities of by-products, making their efficient valorization a pressing challenge [2].This study emphasizes the potential of using polar food-grade solvents, such as water and ethanol, to extract mushroom bioactive compounds with high yields (e.g., Ab-Whole-Aq: 0.56 g/g dw; PO-Stalk-Alk: 0.57 g/g dw), demonstrating their viability as natural food additives. In addition, the insoluble fraction recovered by alkaline extraction, which is partly composed of polysaccharides such as chitin, could be used for the synthesis of chitosan [56] or for food fortification since these extracts are rich in crude fiber and protein [57,58]. Polar solvents, including water and ethanol, have demonstrated their viability in efficiently extracting compounds with functional properties, making these extracts promising candidates for natural food additives, preservatives, and nutraceuticals.

Scaling up these extraction techniques for industrial-scale applications will necessitate taking into account elements like scalability and cost-effectiveness. Even if the extraction methods employed in this study work well in the lab, they may not be suitable for large-scale industrial production because of issues including solvent recovery, energy consumption, and the requirement for high-capacity equipment [59]. Despite these difficulties, the relative ease of usage and the broad availability of food-grade solvents for polar solvent extractions encourage its scalability for industrial use. By promoting circular economy principles and cutting down on waste in the agriculture sector, the use of mushroom by-products in food formulations may offer a viable alternative for the food industry as well as mushroom growers.

In the food sector, mushroom by-products have a variety of uses, mostly as natural preservatives or functional substances [3]. The aqueous extracts’ prebiotic qualities, especially those of *P. ostreatus* and *A. bisporus*, demonstrate their capacity to support gut health by fostering the development of advantageous probiotic bacteria, such *Lactobacillus* strains. Functional meals designed to enhance digestion and the general equilibrium of the gut microbiota may contain these bioactive substances. Additionally, the antibacterial qualities of these extracts, especially in *A. bisporus* and *L. edodes*, imply that they may be used as natural food preservatives, lowering the requirement for artificial chemicals while extending the shelf life of food products. For example, shiitake extracts could be incorporated into meat or dairy products to prevent spoilage caused by pathogenic bacteria, aligning with the growing consumer demand for clean-label, natural ingredients. Furthermore, the antioxidant potential of *A. bisporus* extracts, particularly those high in phenolic compounds, could be used to develop functional foods that protect against oxidative stress, helping to prevent chronic diseases like heart disease, diabetes, and cancer [60]. Incorporating these extracts into food compositions provides both health advantages and adds value to otherwise discarded byproducts.

Despite the promising results of this study, several limitations must be addressed before the industrial application of these extracts. The scalability of the extraction processes is a key concern, as the methods used in this study may not be directly applicable to large-scale production. Scaling up will require optimization of extraction parameters, such as solvent concentration, extraction time, and temperature, to maximize yield while minimizing cost. Additionally, the cost of extraction solvents and energy requirements for large-scale extractions should be evaluated to ensure the economic feasibility of these processes. Another limitation is the variability in the bioactive properties of mushroom by-products, which can vary based on factors such as mushroom species, growth conditions, and post-harvest handling [61]. Future studies should focus on standardizing the production of these by-products and developing methods to ensure consistency in the bioactive compound profiles. This would help ensure reliable performance when used in food applications.

Future research should also look into the possibility of using mushroom by-products into other food industries, such as plant-based meat replacements, functional beverages, and nutraceuticals. Furthermore, the utilization of by-products in the production of packaging materials or biodegradable products is a viable route to pursue in terms of sustainability. The environmental impact of these extraction procedures, including the life cycle analysis of mushroom by-product valorization, will be critical in establishing their sustainability and overall contribution to circular economy practices in agriculture and the food industry.

## 5. Conclusions

This study highlights the potential of mushroom by-products, such as stems and non-commercial mushrooms, as important resources for the food sector, particularly in terms of sustainability and circularity. By the extraction of bioactive compounds with strong prebiotic, antibacterial, and antioxidant effects, this study shows how mushroom by-products can be repurposed to make functional foods that support gut health and extend the shelf life of food products. Extracts high in polysaccharides, such as PO-Stalk-Aq, have been shown to support probiotic growth and improve metabolic activity of probiotic strains, whereas *A. bisporus* and *L. edodes* extracts have antimicrobial and antioxidant properties that highlight their potential as natural food preservatives and additives. Furthermore, the use of polar food-grade solvents such as water and ethanol results in high extraction yields of bioactive compounds, making it an economically viable strategy for incorporating these compounds into food compositions. This research not only helps to develop functional meals, but it also promotes the use of zero-waste methods in the mushroom industry. Following circular economy ideas, valuing mushroom by-products helps to encourage the change toward more environmentally friendly industrial and agricultural methods. Ultimately, this work provides a framework for improving the sustainability, economic efficiency, and environmental impact of mushroom production, offering a model for future research and industrial applications.

## Data availability

The data supporting our findings are available in the manuscript file or from the corresponding author upon request.

## Declaration of competing interest

The authors declare that they have no known competing financial interests or personal relationships that could have appeared to influence the work reported in this paper.

## Funding Sources

This work was supported by grant 230025CONV from the Junta de Comunidades de Castilla-La Mancha (co-financed European Union FEDER funds).

## Ethics statement

Not applicable.

## Author Contributions

Research design and conceptualization: P.N.S, A.R.M., L.G.G. and O.A.. Writing, reviewing, and editing: L.G.G., O.A., P.N.S., A.J.L., and E.M.G.; Polysaccharide extraction and analyses: L.G.G., A. R-M., A.P., and PNS.; Prebiotic and antimicrobial analyses: P.N.S., A.J.L., and E.M.G. All the authors read and approved the manuscript.

## Abbreviations

AB-Whole: *Agaricus Bisporus* – Whole Mushroom
PO-Stalk: *Pleurotus ostreatus* - Stalk
LE-Stem: *Lentinula edodes* - Stem

## Acknowledgements

The authors would like to thank Setas Meli S.L. for mushrooms’ by-products supplies.

## Notes

### Competing Interest Statement

The authors have declared no competing interest.

## References

1. Wan Mahari WA, Peng W, Nam WL, Yang H, Lee XY, Lee YK, et al. A review on valorization of oyster mushroom and waste generated in the mushroom cultivation industry. J Hazard Mater. 2020 Dec 5;400:123156.

2. Guo J, Zhang M, Fang Z. Valorization of mushroom by-products: a review. J Sci Food Agric. 2022;102(13):5593–605.

3. Navarro-Simarro P, Gómez-Gómez L, Ahrazem O, Rubio-Moraga Á. Food and human health applications of edible mushroom by-products. New Biotechnol. 2024 Jul 25;81:43– 56.

4. Abdelshafy AM, Belwal T, Liang Z, Wang L, Li D, Luo Z, et al. A comprehensive review on phenolic compounds from edible mushrooms: Occurrence, biological activity, application and future prospective. Crit Rev Food Sci Nutr. 2022 Jul 23;62(22):6204–24.

5. Golian M, Hegedűsová A, Mezeyová I, Chlebová Z, Hegedűs O, Urminská D, et al. Accumulation of Selected Metal Elements in Fruiting Bodies of Oyster Mushroom. Foods. 2022 Jan;11(1):76.

6. Guan W, Zhang J, Yan R, Shao S, Zhou T, Lei J, et al. Effects of UV-C treatment and cold storage on ergosterol and vitamin D2 contents in different parts of white and brown mushroom (Agaricus bisporus). Food Chem. 20160413th ed. 2016 Nov 1;210:129–34.

7. Ma G, Xu Q, Du H, Muinde Kimatu B, Su A, Yang W, et al. Characterization of polysaccharide from *Pleurotus eryngii* during simulated gastrointestinal digestion and fermentation. Food Chem. 2022 Feb 15;370:131303.

8. Törős G, El-Ramady H, Prokisch J, Velasco F, Llanaj X, Nguyen DHH, et al. Modulation of the Gut Microbiota with Prebiotics and Antimicrobial Agents from Pleurotus ostreatus Mushroom. Foods. 2023 Jan;12(10):2010.

9. Xu X, Yang J, Ning Z, Zhang X. Lentinula edodes-derived polysaccharide rejuvenates mice in terms of immune responses and gut microbiota. Food Funct. 2015;6(8):2653–63.

10. Zhao M, Tang F, Huang X, Ma J, Wang F, Zhang P. Polysaccharide Isolated from Agaricus blazei Murill Alleviates Intestinal Ischemia/Reperfusion Injury through Regulating Gut Microbiota and Mitigating Inflammation in Mice. J Agric Food Chem. 2024 Jan 31;72(4):2202–13.

11. Thikham S, Jeenpitak T, Shoji K, Phongthai S, Therdtatha P, Yawoop A, et al. Pulsed electric field-assisted extraction of mushroom ß-glucan from Pleurotus pulmonarius by-product and study of prebiotic properties. Int J Food Sci Technol. 2024;59(6):3939–49.

12. Erdoğan Eliuz EA. Antibacterial activity and antibacterial mechanism of ethanol extracts of Lentinula edodes (Shiitake) and Agaricus bisporus (button mushroom). Int J Environ Health Res. 2022 Aug 3;32(8):1828–41.

13. Umaña M, Eim V, Garau C, Rosselló C, Simal S. Ultrasound-assisted extraction of ergosterol and antioxidant components from mushroom by-products and the attainment of a β-glucan rich residue. Food Chem. 2020 Dec 1;332:127390.

14. Ramos M, Burgos N, Barnard A, Evans G, Preece J, Graz M, et al. *Agaricus bisporus* and its by-products as a source of valuable extracts and bioactive compounds. Food Chem. 2019 Sep 15;292:176–87.

15. Pérez-Montes A, Rangel-Vargas E, Lorenzo JM, Romero L, Santos EM. Edible mushrooms as a novel trend in the development of healthier meat products. Curr Opin Food Sci. 2021 Feb 1;37:118–24.

16. Vital ACP, Goto PA, Hanai LN, Gomes-da-Costa SM, Filho BAA, Nakamura CV, et al. Microbiological, functional and rheological properties of low fat yogurt supplemented with Pleurotus ostreatus aqueous extract. LWT - Food Sci Technol. 2015;64(2):1028–35.

17. Parvin R, Farzana T, Mohajan S, Rahman H, Rahman SS. Quality improvement of noodles with mushroom fortified and its comparison with local branded noodles. NFS J. 2020 Aug 1;20:37–42.

18. Du J, Xi J, Chen X, Sun H, Zhong L, Zhan Q, et al. Effects of different extraction methods on the release of non-volatile flavor components in shiitake mushroom (*Len4nus edodes*). J Food Compos Anal. 2024 Apr 1;128:106001.

19. Saetang N, Rattanapot T, Manmai N, Amornlerdpison D, Ramaraj R, Unpaprom Y. Effect of hot water extraction process on schizophyllan from split gill mushroom. Biomass Convers Biorefinery. 2024 Jan 1;14(1):1017–26.

20. Huang X, Ai C, Yao H, Zhao C, Xiang C, Hong T, et al. Guideline for the extraction, isolation, purification, and structural characterization of polysaccharides from natural resources. eFood. 2022;3(6):e37.

21. Ghosh S, Nandi S, Banerjee A, Sarkar S, Chakraborty N, Acharya K. Prospecting medicinal properties of Lion’s mane mushroom. J Food Biochem. 2021 Aug;45(8):e13833.

22. Trapero A, Ahrazem O, Rubio-Moraga A, Jimeno ML, Gómez MD, Gómez-Gómez L. Characterization of a Glucosyltransferase Enzyme Involved in the Formation of Kaempferol and Quercetin Sophorosides in Crocus sativus. Plant Physiol. 2012 Aug 1;159(4):1335–54.

23. Yang B, Kortesniemi M, Liu P, Karonen M, Salminen JP. Analysis of Hydrolyzable Tannins and Other Phenolic Compounds in Emblic Leafflower (Phyllanthus emblica L.) Fruits by High Performance Liquid Chromatography–Electrospray Ionization Mass Spectrometry. J Agric Food Chem. 2012 Sep 5;60(35):8672–83.

24. Bermúdez-Gómez P, Fernández-López J, Pérez-Clavijo M, Viuda-Martos M. Evaluation of Sample Size Influence on Chemical Characterization and In Vitro Antioxidant Properties of Flours Obtained from Mushroom Stems Coproducts. Antioxidants. 2024 Mar;13(3):349.

25. Salazar N, Prieto A, Leal JA, Mayo B, Bada-Gancedo JC, de los Reyes-Gavilán CG, et al. Production of exopolysaccharides by *Lactobacillus* and *Bifidobacterium* strains of human origin, and metabolic activity of the producing bacteria in milk. J Dairy Sci. 2009 Sep 1;92(9):4158–68.

26. Abdul Hakim BN, Xuan NJ, Oslan SNH. A Comprehensive Review of Bioactive Compounds from Lactic Acid Bacteria: Potential Functions as Functional Food in Dietetics and the Food Industry. Foods. 2023 Jan;12(15):2850.

27. Ashrafudoulla M, Mevo SIU, Song M, Chowdhury MAH, Shaila S, Kim DH, et al. Antibiofilm mechanism of peppermint essential oil to avert biofilm developed by foodborne and food spoilage pathogens on food contact surfaces. J Food Sci. 2023 Sep;88(9):3935–55.

28. Moreno-Chamba B, Salazar-Bermeo J, Navarro-Simarro P, Narváez-Asensio M, Martinez-Madrid MC, Saura D, et al. Autoinducers modulation as a potential anF-virulence target of bacteria by phenolic compounds. Int J Antimicrob Agents. 2023 Oct 1;62(4):106937.

29. Mondéjar-López M, López-Jimenez AJ, Ahrazem O, Gómez-Gómez L, Niza E. Chitosan coated - biogenic silver nanoparticles from wheat residues as green antifungal and nanoprimig in wheat seeds. Int J Biol Macromol. 2023 Jan 15;225:964–73.

30. Jiménez-Zamora A, Pastoriza S, Rufián-Henares JA. Revalorization of coffee by-products. Prebiotic, antimicrobial and antioxidant properties. LWT - Food Sci Technol. 2015 Apr 1;61(1):12–8.

31. Borshchevskaya LN, Gordeeva TL, Kalinina AN, Sineokii SP. Spectrophotometric determination of lactic acid. J Anal Chem. 2016 Aug 1;71(8):755–8.

32. Lin X, Sun DW. Research advances in browning of button mushroom (*Agaricus bisporus*): Affecting factors and controlling methods. Trends Food Sci Technol. 2019 Aug 1;90:63–75.

33. Souza D de, Sbardelotto AF, Ziegler DR, Marczak LDF, Tessaro IC. Characterization of rice starch and protein obtained by a fast alkaline extraction method. Food Chem. 2016 Jan 15;191:36–44.

34. Ojeaburu SI, Oriakhi K. Hepatoprotective, antioxidant and, anF-inflammatory potentials of gallic acid in carbon tetrachloride-induced hepatic damage in Wistar rats. Toxicol Rep. 20210108th ed. 2021;8:177–85.

35. Gutierrez-Rodelo C, Ochoa-Lopez A, Luis Balderas-Lopez J, Reyes-Ramirez A, Millan-Pacheco C, Favela-Rosales F, et al. Eritadenine as a regulator of anxiety Disorders: An experimental and docking Approach. Neurosci Lett. 20230803rd ed. 2023 Sep 14;813:137413.

36. Ukaegbu CI, Shah SR, Hamid HA, Alara OR, Sarker MdZI. Phenolic Compounds of Aqueous and Methanol Extracts of Hypsizygus tessellatus (brown and white var.) and Flammulina velutipes caps: Antioxidant and Antiproliferative Activities. Pharm Chem J. 2020 May 1;54(2):170–83.

37. Ahmad I, Arif M, Xu M, Zhang J, Ding Y, Lyu F. Therapeutic values and nutraceutical properties of shiitake mushroom (Lentinula edodes): A review. Trends Food Sci Technol. 2023 Apr 1;134:123–35.

38. Garcia J, Afonso A, Fernandes C, Nunes FM, Marques G, Saavedra MJ. Comparative antioxidant and antimicrobial properties of *Len4nula edodes* Donko and Koshin varieties against priority multidrug-resistant pathogens. South Afr J Chem Eng. 2021 Jan 1;35:98– 106.

39. Zeng C, Ye G, Li G, Cao H, Wang Z, Ji S. RID serve as a more appropriate measure than phenol sulfuric acid method for natural water-soluble polysaccharides quantication. Carbohydr Polym. 2022 Feb 15;278:118928.

40. Aranaz I, Alcántara AR, Civera MC, Arias C, Elorza B, Heras Caballero A, et al. Chitosan: An Overview of Its Properties and Applications. Polymers. 2021 Jan;13(19):3256.

41. Rastogi S, Singh A. Gut microbiome and human health: Exploring how the probiotic genus Lactobacillus modulate immune responses. Front Pharmacol. 2022;13:1042189.

42. Chee WJY, Chew SY, Than LTL. Vaginal microbiota and the potential of Lactobacillus derivatives in maintaining vaginal health. Microb Cell Factories. 2020 Nov 7;19(1):203.

43. Wang Y, Wu J, Lv M, Shao Z, Hungwe M, Wang J, et al. Metabolism Characteristics of Lactic Acid Bacteria and the Expanding Applications in Food Industry. Front Bioeng Biotechnol. 2021;9:612285.

44. Karnaouri A, Asimakopoulou G, Kalogiannis KG, Lappas A, Topakas E. Efficient d-lactic acid production by *Lactobacillus delbrueckii* subsp. *bulgaricus* through conversion of organosolv pretreated lignocellulosic biomass. Biomass Bioenergy. 2020 Sep 1;140:105672.

45. Sawangwan T, Wansanit W, Pattani L, Noysang C. Study of prebiotic properties from edible mushroom extraction. Agric Nat Resour. 2018;52(6):519–24.

46. Nowak R, Nowacka-Jechalke N, Juda M, Malm A. The preliminary study of prebiotic potential of Polish wild mushroom polysaccharides: the stimulation effect on Lactobacillus strains growth. Eur J Nutr. 20170328th ed. 2017 Jun;57(4):1511–21.

47. Kim JH, Tam CC, Chan KL, Mahoney N, Cheng LW, Friedman M, et al. Antimicrobial Efficacy of Edible Mushroom Extracts: Assessment of Fungal Resistance. Appl Sci. 2022 Jan;12(9):4591.

48. Moussa AY, Fayez S, Xiao H, Xu B. New insights into antimicrobial and antibiofilm effects of edible mushrooms. Food Res Int. 2022 Dec 1;162:111982.

49. Yadav D, Prashanth KVH, Negi PS. Low molecular weight chitosan from *Pleurotus ostreatus* waste and its prebiotic potential. Int J Biol Macromol. 2024 May 1;267:131419.

50. Fukushima-Sakuno E. Bioactive small secondary metabolites from the mushrooms Lentinula edodes and Flammulina velutipes. J Antibiot (Tokyo). 2020 Oct;73(10):687–96.

51. Kong Y, Zhang LL, Zhao J, Zhang YY, Sun BG, Chen HT. Isolation and identication of the umami peptides from shiitake mushroom by consecutive chromatography and LC-Q-TOF-MS. Food Res Int. 20181129th ed. 2019 Jul;121:463–70.

52. Hutkins R, Walter J, Gibson GR, Bedu-Ferrari C, Scott K, Tancredi DJ, et al. Classifying compounds as prebiotis — scientific perspectives and recommendations. Nat Rev Gastroenterol Hepatol. 2024 Oct 2;1–17.

53. Wang L, Guo H, Liu X, Jiang G, Li C, Li X, et al. Roles of Lentinula edodes as the pork lean meat replacer in production of the sausage. Meat Sci. 20190517th ed. 2019 Oct;156:44– 51.

54. Radzki W, Tutaj K, Skrzypczak K, Michalak-Majewska M, Gustaw W. Ethanolic Extracts of Six Cultivated Mushrooms as a Source of Bioactive Compounds. Appl Sci. 2024 Jan;14(1):66.

55. Nitthikan N, Leelapornpisid P, Naksuriya O, Intasai N, Kiapsin K. Potential and Alternative Bioactive Compounds from Brown Agaricus bisporus Mushroom Extracts for Xerosis Treatment. Sci Pharm. 2022 Dec;90(4):59.

56. Alimi BA, Pathania S, Wilson J, Duffy B, Frias JMC. Extraction, quantification, characterization, and application in food packaging of chitin and chitosan from mushrooms: A review. Int J Biol Macromol. 2023 May 15;237:124195.

57. Zhang Y, Zhu J, Zou Y, Ye Z, Guo L, Zheng Q. Insoluble dietary fiber from five commercially cultivated edible mushrooms: Structural, physiochemical and functional properties. Food Biosci. 2024 Feb 1;57:103514.

58. González A, Nobre C, Simões LS, Cruz M, Loredo A, Rodríguez-Jasso RM, et al. Evaluation of functional and nutritional potential of a protein concentrate from *Pleurotus ostreatus* mushroom. Food Chem. 2021 Jun 1;346:128884.

59. Picot-Allain C, Mahomoodally MF, Ak G, Zengin G. Conventional versus green extraction techniques — a comparative perspective. Curr Opin Food Sci. 2021 Aug 1;40:144–56.

60. Chen X, Li H, Zhang B, Deng Z. The synergistic and antagonistic antioxidant interactions of dietary phytochemical combinations. Crit Rev Food Sci Nutr. 2022 Jul 8;62(20):5658–77.

61. Assemie A, Abaya G. The Effect of Edible Mushroom on Health and Their Biochemistry. Int J Microbiol. 2022;2022(1):8744788.

